# Anchorage of H3K9-methylated heterochromatin to the nuclear periphery helps mediate P-cell nuclear migration though constricted spaces in *Caenorhabditis elegans*

**DOI:** 10.1101/2024.05.22.595380

**Authors:** Ellen F. Gregory, G.W. Gant Luxton, Daniel A. Starr

## Abstract

Nuclei adjust their deformability while migrating through constrictions to enable structural changes and maintain nuclear integrity. The effect of heterochromatin anchored at the nucleoplasmic face of the inner nuclear membrane on nuclear morphology and deformability during *in vivo* nuclear migration through constricted spaces remains unclear. Here, we show that abolishing peripheral heterochromatin anchorage by eliminating CEC-4, a chromodomain protein that tethers H3K9-methylated chromatin to the nuclear periphery, disrupts constrained P-cell nuclear migration in *Caenorhabditis elegans* larvae in the absence of the established LINC complex-dependent pathway. CEC-4 acts in parallel to an actin and CDC-42-based pathway. We also demonstrate the necessity for the chromatin methyltransferases MET-2 and JMJD-1.2 during P-cell nuclear migration in the absence of functional LINC complexes. We conclude that H3K9-nethylated chromatin needs to be anchored to the nucleoplasmic face of the inner nuclear membrane to help facilitate nuclear migration through constricted spaces *in vivo*.

## INTRODUCTION

Migrating cells must squeeze through constrictions as part of developmental and pathological processes, including immune cell extravasation and cancer metastasis (McGregor *et al*. 2016; Paul *et al*. 2017). Deformation of the nucleus, the largest and stiffest organelle, is often the rate-limiting step of cell migration through tight spaces (Davidson *et al*. 2014; Thiam *et al*. 2016; Renkawitz *et al*. 2019). Most studies of nuclear migration through constrictions are limited to *in vitro* models (Raab *et al*. 2016; Denais *et al*. 2016; Bone and Starr 2016; Xia *et al*. 2019). Here we use an *in vivo* model, where *C. elegans* P-cell nuclei migrate through a constricted space as a normal part of larval development (Chang *et al*. 2013; Bone and Starr 2016; Bone *et al*. 2016). During this process, nuclei of neuronal and vulval precursor cells, referred to as P cells, move from lateral positions to the ventral cord of the organism (Sulston and Horvitz 1977; Bone *et al*. 2016; Fridolfsson *et al*. 2018). To complete this migration, P-cell nuclei must pass through a constriction between the cuticle and body wall muscle that is 100-200 nm, ∼5% of the diameter of P-cell nuclei prior to migration (Bone *et al*. 2016). Following nuclear migration to the ventral cord, P cells divide and differentiate to form the vulva and twelve of nineteen gamma-aminobutyric acid (GABA)-producing (GABAergic) neurons (Sulston and Horvitz 1977).

Forward genetic screens for mutants missing P-cell derived lineages have identified three parallel pathways that move P-cell nuclei through constrictions. The first pathway is the linker of nucleoskeleton and cytoskeleton (LINC) complex pathway, which includes the Klarsicht/ANC-1/SYNE homology (KASH) protein UNC-83 in the outer nuclear membrane and the Sad1/UNC-84 (SUN) protein UNC-84 in the inner nuclear membrane (Malone *et al*. 1999; Starr *et al*. 2001; Starr 2019). UNC-83 and UNC-84 interact with one another in the perinuclear space to create a force-transducing bridge across the nuclear envelope (McGee *et al*. 2006). UNC-84 interacts with lamins in the nucleoplasm, while the cytoplasmic domain of UNC-83 recruits cytoplasmic dynein to the outer nuclear membrane to couple the nucleoskeleton to the microtubule cytoskeleton (Fridolfsson *et al*. 2010; Bone *et al*. 2014; Gregory *et al*. 2023). Dynein then generates the forces to move P-cell nuclei toward the minus ends of microtubules (Bone *et al*. 2016; Ho *et al*. 2018). The second pathway is an actin-based pathway that involves the small GTPase cell division control protein 42 (CDC-42) (Chang *et al*. 2013) and CDC-42 guanine nucleotide exchange factor-1 (CGEF-1), a predicted GEF for CDC-42 (Ho *et al*. 2023). In addition, auxin-induced degradation of CDC-42 or the actin-branching actin-related protein 2/3 (Arp2/3) complex subunit ARX-3 in larval P cells also induced strong nuclear migration defects, suggesting that these actin regulators work together in a common pathway to help move P-cell nuclei through constrictions (Ho *et al*. 2023). In the third pathway, mutations in filamin-2 (*fln-2*) led to an increase in nuclear rupturing, suggesting FLN-2 functions to maintain and/or repair the nuclear envelope during P-cell nuclear migration (Ma *et al*. 2023).

Inhibiting any of the LINC complex, actin/CDC-42, or FLN-2-dependent pathways on their own leads to no or mild nuclear migration defects (Chang *et al*. 2013; Bone *et al*. 2016; Ho *et al*. 2023; Ma *et al*. 2023). Moreover, simultaneously inhibiting all three pathways lead to a severe, but still incompletely penetrant phenotype (Ma *et al*. 2023). Therefore, we hypothesized that another pathway functions in parallel to these three previously identified pathways to help move P-cell nuclei through constricted spaces.

In other systems, heterochromatin is thought to safeguard nuclear integrity from mechanical stresses during nuclear movements and shape changes (Kalukula *et al*. 2022). Heterochromatin facilitates mammalian tissue culture cell migration through micropores in a transcription-independent manner (Gerlitz and Bustin 2010; Krause *et al*. 2019; Hsia *et al*. 2022). In a similar manner, inhibiting histone H3K4 methyltransferases affected the physical properties of nuclei and abrogated interstitial T cell migration through a three-dimensional matrix as well as in a zebrafish xenotransplantation model (Wang *et al*. 2018). The peripheral localization of heterochromatin at the nucleoplasmic face of the inner nuclear membrane appears to be critical for its function in regulating nuclear dynamics (Penagos-Puig and Furlan-Magaril 2020). Untethering heterochromatin from the nuclear envelope increases nuclear deformability in the fission yeast *Schizosaccharomyces pombe* (Schreiner *et al*. 2015). Based on these previously reported findings, we explored how peripheral heterochromatin might function independently of gene expression during *C. elegans* P-cell nuclear migration. Here, we identify a fourth pathway where the inner nuclear membrane protein CEC-4, which anchors heterochromatin at the nuclear periphery (Gonzalez-Sandoval *et al*. 2015), facilitates larval P-cell nuclear migration through constricted spaces.

## RESULTS and DISCUSSION

### Untethering H3K9-methylated heterochromatin from the inner nuclear membrane contributes to P-cell nuclear migration failure

We hypothesized that heterochromatin anchored at the nuclear periphery affects nuclear deformability, thus impacting the capability of P-cell nuclei to migrate through narrow spaces. To test this hypothesis, we asked if P-cell nuclear migration occurred normally in *cec-4* null animals. The *C. elegans* inner nuclear membrane protein CEC-4 anchors H3K9-methylated chromatin to the periphery of the nuclear envelope (Gonzalez-Sandoval *et al*. 2015). Mutating *cec-4* untethers H3K9-methylated heterochromatin from the nucleoplasmic face of the inner nuclear membrane without affecting global transcription (Gonzalez-Sandoval *et al*. 2015). This approach enabled us to assess the influence of inner nuclear membrane-anchored heterochromatin on *in vivo* nuclear migration independently of any alterations in gene expression.

We assay P-cell nuclear migration by scoring missing cells in the P-cell lineage after nuclei move to the ventral cord (Fridolfsson *et al*. 2018). Normally, 12 of 19 GABAergic neurons are derived from P cells. Thus, defects in P-cell nuclear migration are expected to have significantly fewer GABAergic neurons than wild-type animals. *cec-4(ok3124)* null mutant animals had an average of 19 GABAergic neurons at 15, 20, and 25° C, indicating that untethering H3K9-methylated heterochromatin from the nuclear periphery, on its own, does not disrupt P-cell nuclear migration (Figure 1). We then crossed the *cec-4* mutant into the sensitized, LINC complex-defective *unc-84(n369)* null background. *unc-84(null)* mutants are missing 3-5 GABAergic neurons at higher temperatures, and are nearly wild type at 15°C. In *unc-84(n369)* animals, loss of *cec-4* significantly increased the number of missing GABA neurons relative to controls. These phenotypes were comparable to those observed in the *cgef-1(yc3) unc-84(n369)* double mutant (Ho *et al*. 2023), in which both the LINC complex and actin/CDC-42-dependent P-cell nuclear migration pathways are disrupted. We conclude that the anchorage of heterochromatin to the nucleoplasmic face of the inner nuclear membrane is critical for P-cell nuclear migration in LINC complex-defective larvae.

**Figure 1:**
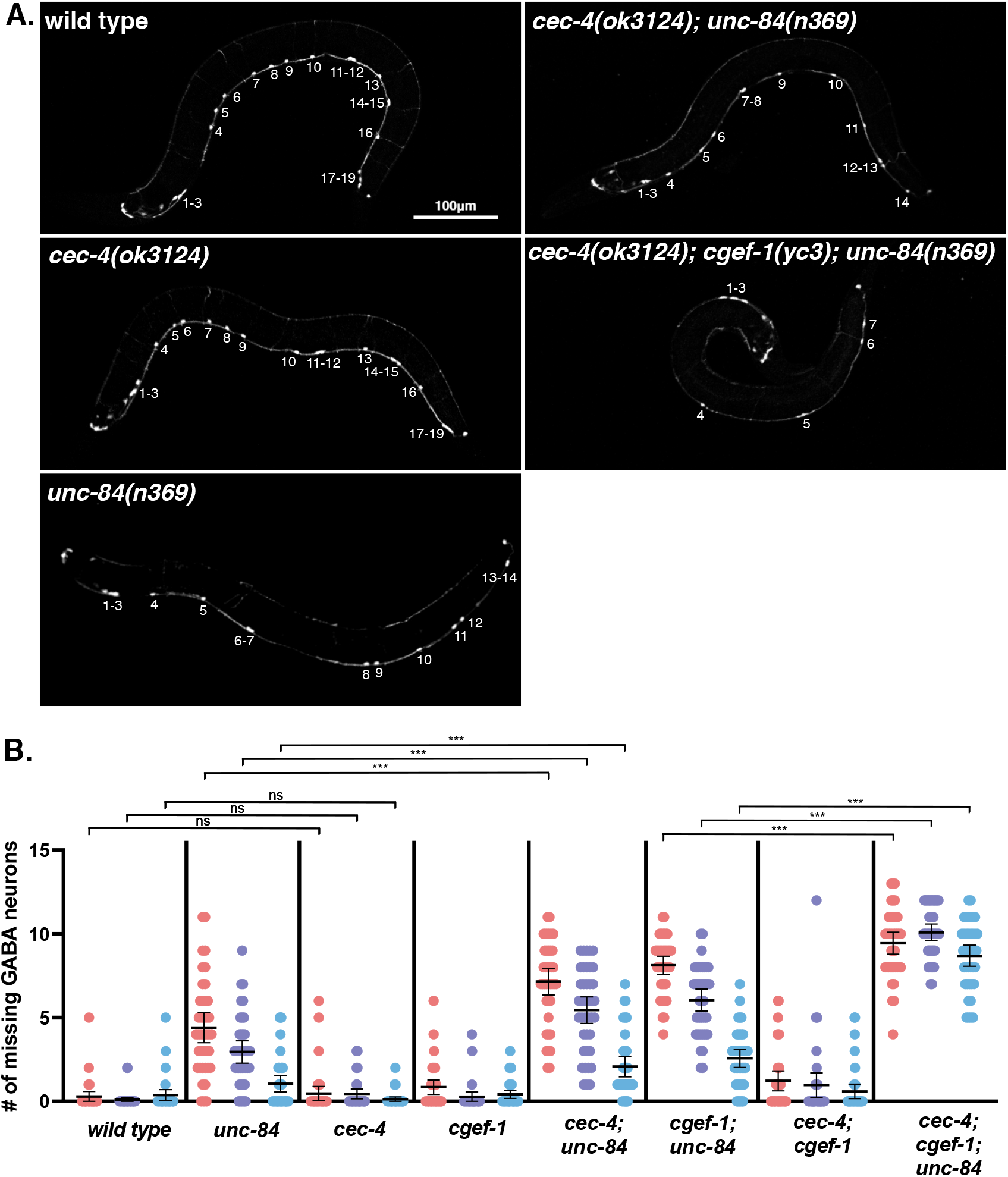
The *cec-4* null mutation enhances the P-cell nuclear migration defect observed in *unc-84* and *unc-84; cgef-1* null mutants. **A)** Images of L4 animals expressing GFP from the *oxIs12[p*_*unc-47*_*::gfp]* marker in P-cell derived GABAergic neurons. Dorsal is up, ventral is down. The numbers of GABAergic neurons present in the ventral cord were scored and are numbered from anterior (left) to posterior. Scale bar: 100 µm. **B)** Plot of the number of missing GABAergic neurons from the 19 observed in *wild type* (UD87 RFP+ animals) and null mutant lines *unc-84* (UD87 RFP-), *cec-4* (UD462 RFP+), *cgef-1* (UD285 RFP+), *cec-4; unc-84* (UD462 RFP-), *cgef-1, unc-84* (UD285 RFP-), *cec-4; cgef-1* (UD795 RFP+), *cec-4; cgef-1, unc-84* (UD795 RFP-). Each dot represents a single worm. Three different temperatures are shown: 25° C (salmon), 20° C (violet), and 15° C (blue). n=40. Means and 95% confidence intervals (CI) are shown. Tests for statistical significance are shown to compare *cec-4* to *wild type, cec-4; unc-84* double mutants to *unc-84* single mutants, and *cec-4; cgef-1, unc-84* triple mutants to *cgef-1, unc-84* double mutants. Results were corrected using a Tukey HSD test. ***p≤0.001

We next simultaneously disrupted all three pathways (i.e., the LINC complex-, the actin/CDC-42-, and the CEC*-*4-dependent pathways). The triple *cec-4; cgef-1, unc-84* mutant had a nearly complete nuclear migration failure at 15, 20, and 25° C (Figure 1). These results suggest that CGEF-1 and peripheral heterochromatin act redundantly to suppress the *unc-84(n369)* nuclear migration defect.

### Depleting H3K9-methylated heterochromatin in LINC complex-defective animals enhanced defects in P-cell nuclear migration

*C. elegans* encodes a pair of H3K9 methyltransferases, MET-2 and SET-25 (Ahringer and Gasser 2018). MET-2, a homolog of mammalian SET domain bifurcated histone lysine methyltransferase 1 (SETDB1), is the primary methyltransferase for H3K9me1 and me2 (Towbin *et al*. 2012). SET-25 plays a secondary role in H3K9me1 and me2 but is the only methyl transferase for H3K9me3 (Towbin *et al*. 2012). Furthermore, *met-2(n4256); set-25(n5021)* double mutant animals exhibit disrupted anchorage of heterochromatin to the nuclear periphery resulting in a phenotype similar to what was observed in *cec-4(ok3124)* mutant animals (Towbin *et al*. 2012; Gonzalez-Sandoval *et al*. 2015). We focused the remainder of our genetic analyses on *met-2*, which encodes the primary H3K9 methyltransferase.

Specifically, we next tested whether knocking out *met-2* in a *cgef-1(yc3) unc-84(n369)* mutant background would phenocopy the P-cell nuclear migration defect observed in the *cec-4(ok3124); cgef-1(yc3) unc-84(n369)* triple null mutant. To do so, we crossed the predicted null allele *met-2(n4256)* into *cgef-1(yc3)* and *unc-84(n369)* single mutants, and *cgef-1(yc3) unc-84(n369)* double mutants and quantified P-cell nuclear migration at 15, 20, and 25° C. Animals with the *met-2(n4256)* mutation in a *cgef-1(yc3)* background had the normal number of GABAergic neurons (Figure 2). However, the presence of the *met-2(null)* allele enhanced the number of missing GABAergic neurons in an *unc-84(n369)* mutant (Figure 2). *met-2(n4256); unc-84(null)* double mutant animals had a similar, perhaps slightly less severe, P-cell nuclear migration defect compared to the phenotypes found in *cgef-1 unc-84* double mutants (Ho *et al*. 2023) (Figure 2). Thus, *met-2* and *unc-84* mutants are synergistic, leading us to conclude that *met-2* functions to help P-cell nuclei migrate in a pathway that operates in parallel to the LINC complex-dependent pathway.

**Figure 2:**
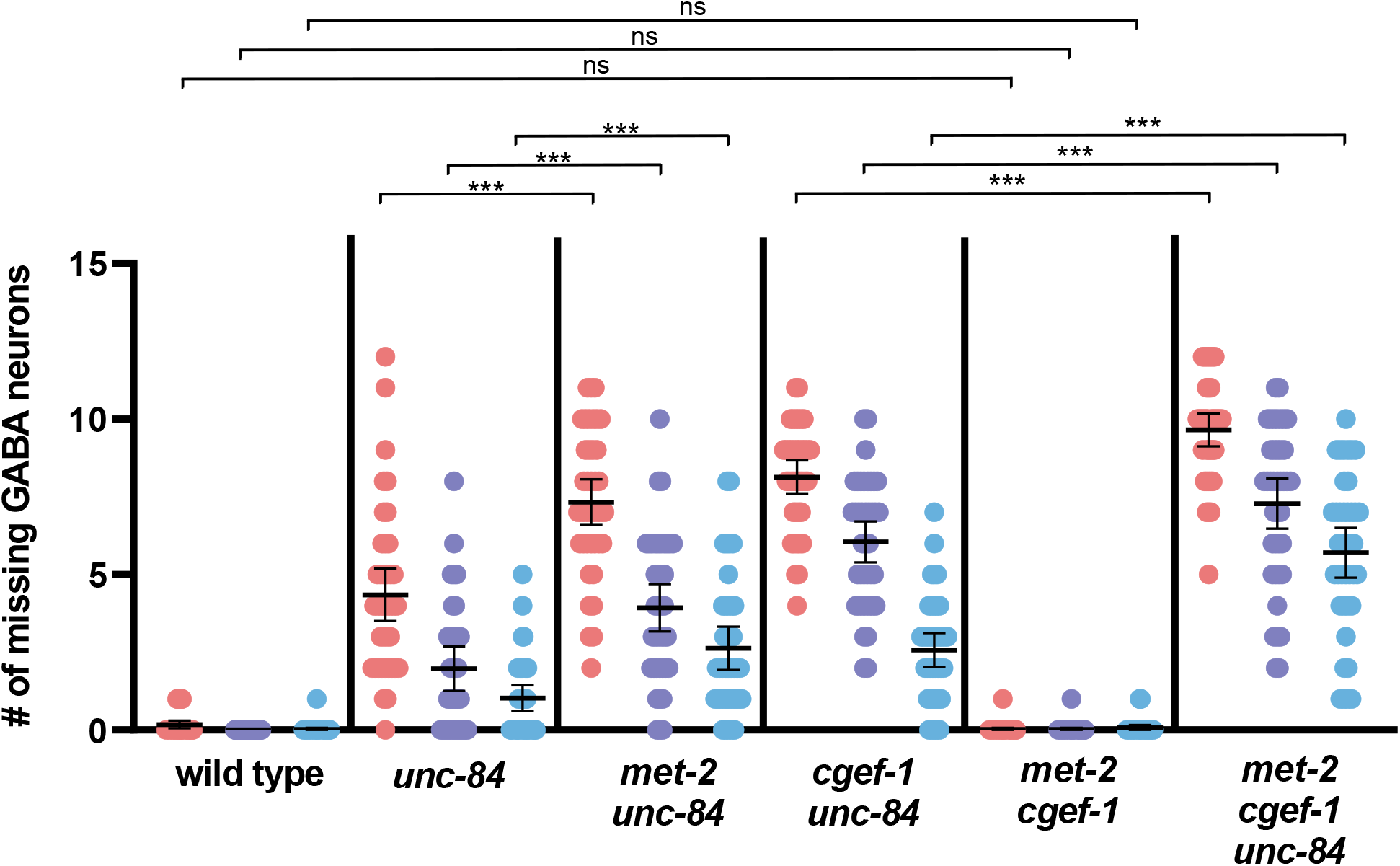
Inhibiting H3K9 methylation enhances the P-cell nuclear migration defects observed in *unc-84* mutants. Plot of the number of missing GABAergic neurons observed in *wild type* (UD87 RFP+ animals) and null mutant lines *unc-84* (UD87 RFP-), *met-2; unc-84* (UD906 RFP-), *cgef-1, unc-84* (UD285 RFP-), *met-2; cgef-1* (UD957 RFP+), *met-2; cgef-1, unc-84* (UD957 RFP-). Each dot represents a single worm. Three different temperatures are shown: 25° C (salmon), 20° C (violet), and 15° C (blue). n=40. Means and 95% CI are shown. Significance indicated is compared to *unc-84(n369)* at each temperature. Results were corrected using a Tukey HSD test. ***p≤0.001.

We then asked whether *met-2* represents a pathway distinct from the actin/CDC-42- and LINC complex-dependent pathways. *met-2(n4256); cgef-1(yc3) unc-84(n369)* triple mutant animals had significantly enhanced P-cell nuclear migration defects compared to those observed in double mutant animals (Figure 2). In fact, at higher temperatures, the triple mutant phenotype approached the theoretical phenotypic maximum of 12 missing GABAergic neurons, suggesting a nearly completely penetrant P-cell nuclear migration defect.

### Inhibiting the JMJD-1.2 demethylase in the *unc-84(null)* background leads to P-cell nuclear migration defects

Decompaction of heterochromatin induced by histone methyltransferase or deacetylase inhibitors alters nuclear structure and inhibits cell migration (Gerlitz and Bustin 2010; Stephens *et al*. 2018). Likewise, treatment with histone demethylase inhibitors can rescue abnormal nuclear morphology even in the presence of disrupted lamins (Stephens *et al*. 2018). Thus, we hypothesized that knocking down a H3K9 demethylase would suppress the P-cell nuclear migration defect of *unc-84; cgef-1* null mutants. To test this, we investigated the role of *jmjd-1*.*2*, which encodes a JmjC demethylase that targets H3K27me2 and H3K9me2 repressive marks (Kleine-Kohlbrecher *et al*. 2010), during P-cell nuclear migration. We used a heterozygous *jmjd-1*.*2(ok3628)* balanced line, referred to as *jmjd-1*.*2(ok3628)/+*, because the homozygous *jmjd-1*.*2* animals are embryonic lethal (Myers *et al*. 2018).

*jmjd-1*.*2/+* animals in an otherwise wild-type background did not exhibit a P-cell nuclear migration defect (Figure 3). However, the *jmjd-1*.*2(ok3268)/+* mutation significantly enhanced the *unc-84(n369)* P-cell nuclear migration defects at 20°C and 25°C (Figure 3). Furthermore, *jmjd-1*.*2(ok3268)/+; cgef-1(yc3) unc-84(n369)* triple mutant animals had significantly more missing GABAergic neurons than the double mutants, approaching the maximum phenotype of 12 expected if they fully inhibited P-cell nuclear migration (Figure 3). In conclusion, the elimination of one copy of the *jmjd-1*.*2*, a gene that encodes for a demethylase, resulted in a disruption of P-cell nuclear migration, resembling the pattern observed in *met-* 2-null animals, which are known to impede most H3K9 methylation.

**Figure 3:**
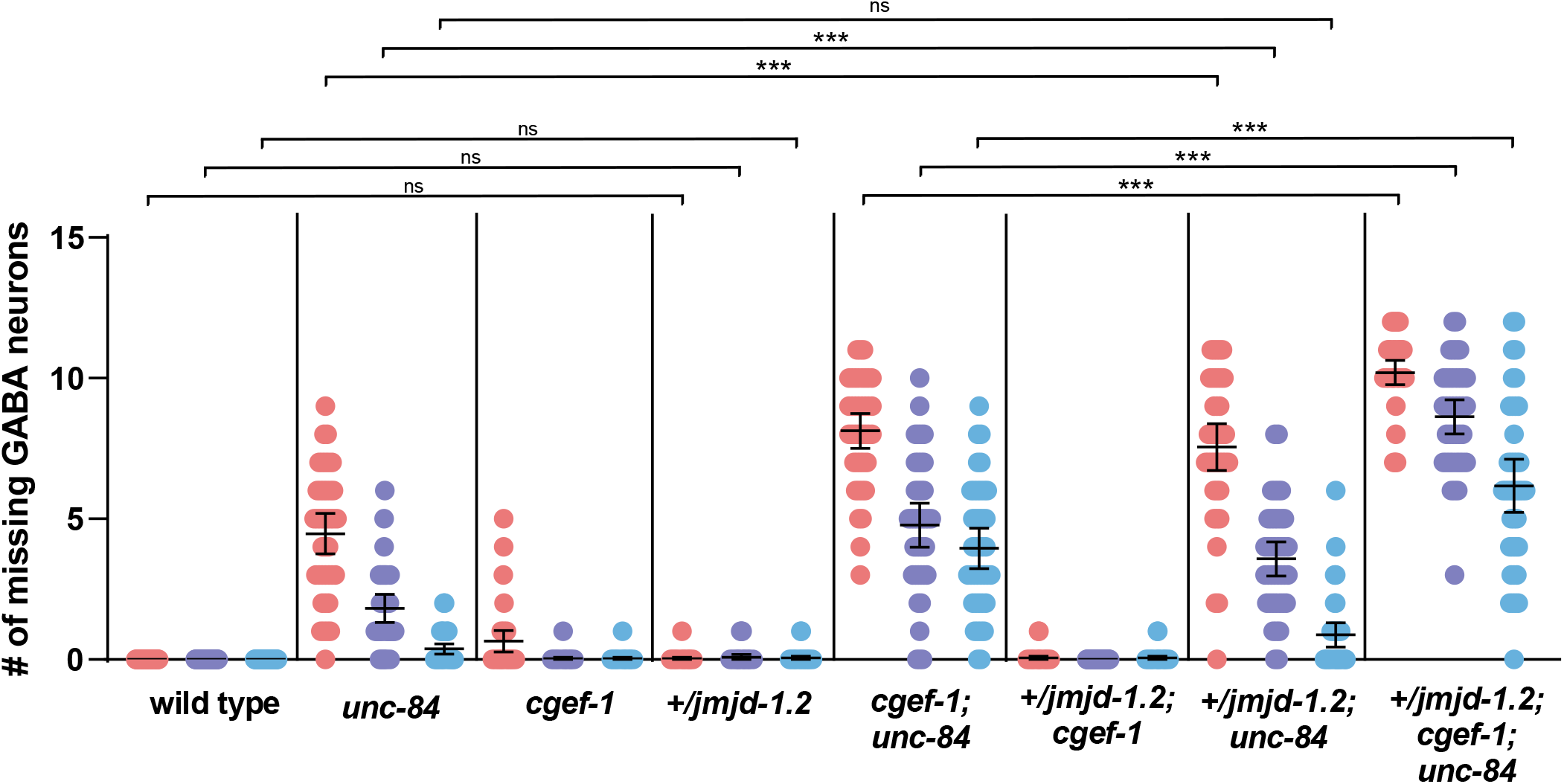
The presence of one copy of the *jmjd-1*.*2* mutation enhances the P-cell nuclear migration observed in *unc-84* mutant animals. A plot of the number of missing GABAergic neurons of *wild type* (UD87 RFP+ animals) and the following null mutant lines: *unc-84* (UD87 RFP-), *cgef-1* (UD285 RFP+), *+/jmjd-1*.*2* (UD996 RFP+) *cgef-1, unc-84* (UD285 RFP-), *+/jmjd-1*.*2; unc-84* (UD996 RFP-), *+/jmjd-1*.*2; cgef-1* (UD997 RFP+), *+/jmjd-1*.*2; cgef-1, unc-84* (UD997 RFP-). Each dot represents a single worm. Three different temperatures are shown: 25° C (salmon), 20° C (violet), and 15° C (blue). n=40. Means and 95% CI are shown. Significance indicated is compared to *unc-84(n369)* at each temperature. Results were corrected using a Tukey HSD test. ***p≤0.001.

## Conclusions

Our genetic data support a model where the anchorage of H3K9-methylated heterochromatin at the nucleoplasmic face of the inner nuclear membrane via CEC-4 plays an important role in nuclear migration through constricted spaces as a part of normal *C. elegans* development. This CEC-4/heterochromatin nuclear anchorage-based pathway works in synergy with three other pathways: the LINC complex pathway (Bone *et al*. 2016), an actin/CDC-42-based pathway (Ho *et al*. 2023), and a FLN-2 based pathway (Ma *et al*. 2023).

It remains unclear how anchoring heterochromatin to the inner nuclear membrane is related to the function of LINC complexes. One possibility could be that LINC complexes transmit mechanical strain to the peripheral heterochromatin (Kirby and Lammerding 2018; Lityagina and Dobreva 2021; Kalukula *et al*. 2022). LINC complexes are required to stabilize the architecture of the nuclear envelope in cells subjected to increased mechanical forces (Crisp *et al*. 2006; Cain *et al*. 2014). Heterochromatin has been implicated in regulating nuclear mechanics (Stephens *et al*. 2019), suggesting that the absence of peripheral heterochromatin could lead to decreased nuclear morphological changes and/or increased nuclear envelope rupturing (Starr 2012; Maciejowski and Hatch 2020). Disruption of LINC complexes upregulates the deposition of heterochromatic marks in *Drosophila* muscle nuclei (Pavlov *et al*. 2023). Alternatively, LINC may regulate the distribution or localization of MET-2 phase-separated foci at the inner nuclear envelope. MET-2 foci rely on CEC-4 for localization at the nuclear periphery, but elimination of CEC-4 does not affect foci formation or number (Delaney *et al*. 2019). However, MET-2 foci disperse in response to higher temperature, resulting in germline developmental defects (Delaney *et al*. 2019). This could be related to the temperature-sensitive P-cell nuclear migration defects seen in LINC null mutant animals (Malone *et al*. 1999; Starr *et al*. 2001). We propose that LINC complexes regulate nuclear mechanics by modulating perinuclear heterochromatin. Since CEC-4 is necessary for heterochromatin localization but not transcriptional repression (Gonzalez-Sandoval *et al*. 2015), our findings likely reflect the requirement for changes in heterochromatin organization rather than transcription during P-cell nuclei migration.

## MATERIALS AND METHODS

### *C. elegans* strains and genetics

Animals were grown on nematode growth medium plates spotted with OP50 *Escherichia coli* (Brenner 1974). Strains were maintained at room temperature (22-23° C), other than those carrying *met-2* and *set-25* mutations, which were kept at 15° C to avoid progressive germline sterility at higher temperatures (Zeller *et al*. 2016). The strains made and used in this paper are listed in Table 1.

**Table 1.** Strains used in this study.

| Strain | Genotype | Reference |
| --- | --- | --- |
| EG1285 | *oxIs12[p<sub>unc-47</sub>::gfp] X* | (McIntire *et al.* 1997) |
| UD87 | *unc-84(n369), oxIs12 X; ycEx60[odr-1::rfp, WRM0617cH07 unc-84(+)]* | (Chang *et al.* 2013) |
| RB2301 | *cec-4(ok3124) IV* | (Consortium 2012) |
| UD642 | *cec-4(ok3124) IV; unc-84(n369), oxIs12 X; ycEx60* | This study |
| UD285 | *cgef-1(yc3), unc-84(n369), oxIs12 X; ycEx60* | (Chang *et al.* 2013) |
| UD795 | *cec-4(ok3124) IV; cgef-1(yc3), unc-84(n369), oxIs12 X; ycEx60* | This study |
| MT13293 | *met-2(n4256) III* | (Lee *et al.* 2019) |
| UD906 | *met-2(n4256) III; unc-84(n369), oxIs12 X; ycEx60* | This study |
| UD957 | *met-2(n4256) III; cgef-1(yc3), unc-84(n369), oxIs12 X; ycEx60* | This study |
| VC2878 | *jmjd-1.2(ok3628) IV / nT1[qIs51] (IV;V)* | (Consortium 2012) |
| UD996 | *jmjd-1.2(ok3628) IV / nT1[qIs51] (IV;V); unc-84(n369), oxIs12 X; ycEx60* | This study |
| UD997 | *jmjd-1.2(ok3628) IV / nT1[qIs51] (IV;V); cgef-1(yc3), unc-84(n369), oxIs12 X; ycEx60* | This study |

### P-cell nuclear migration assay

Successful nuclear migration of larval P cells was scored by counting the number of GABAergic neurons marked with *oxIs12[p*_*unc-47*_::*gfp]* (McIntire *et al*. 1997) in animals at the fourth larval stage of development, as described (Fridolfsson *et al*. 2018). GABAergic neurons and the *p*_*odr-1*_*::RFP* marker were visualized using a wide-field epifluorescent Leica DM6000 microscope with a 63 × Plan Apo 1.40 NA objective, a Leica DC350 FX camera, and Leica LAS AF software (Leica Microsystems, Inc., Deerfield, IL).

### Statistics

Data were tested for significance at each temperature (15°C, 20°C, and 25°C) with one-way ANOVA. Results were corrected by Tukey honestly significant difference (HSD) test calculated using the R multcomp package v1.4-23 (Hothorn *et al*. 2008). Graphs were made using Prism 9 software (GraphPad, Boston, MA).

## Acknowledgements

We thank past and present members of the Starr-Luxton lab for their input on this research. We thank the *Caenorhabditis* Genetics Center (CGC), which is funded by the National Institutes of Health Office of Research Infrastructure Programs (P40OD010440), for providing strains. We also thank WormBase.

## Author contributions

Conceptualization: E.F.G., G.W.G.L., and D.A.S.; Methodology: E.F.G. and D.A.S.; Validation: E.F.G.; Formal analysis: E.F.G.; Investigation: E.F.G.; Resources: E.F.G., G.W.G.L., and D.A.S.; Data curation: E.F.G.; Writing – original draft: E.F.G. and D.A.S.; Writing – review and editing: E.F.G., G.W.G.L., and D.A.S.; Visualization: E.F.G. and D.A.S.; Supervision: G.W.G.L. and D.A.S.; Project administration: D.A.S.; Funding acquisition: D.A.S.

## Data availability

Strains are available upon request. The authors affirm that all data necessary for confirming the conclusions of the article are present within the article, figures, and tables.

